# Distinct regimes of spatial prediction across the visual field during natural vision

**DOI:** 10.64898/2026.03.27.714859

**Authors:** Wieger H. Scheurer, Micha Heilbron

## Abstract

Prediction is foundational to theories of perceptual processing and learning, in both neuroscience and AI. However, it remains unclear whether prediction occurs routinely during naturalistic perception, and at what level of abstraction the brain predicts. Here, we address both questions by analysing 7T fMRI recordings of humans viewing 73,000 natural images. We use deep generative models to quantify spatial predictability at multiple levels of abstraction, and relate these to retinotopically precise responses across V1-V4, while rigorously controlling for local image features. This reveals that, even during natural scene viewing, responses throughout visual cortex are modulated by spatial predictability, with more predictable inputs evoking weaker responses. In central vision, we observe a hierarchy of predictions that parallels the feature-encoding gradient: V1 is most sensitive to low-level unpredictability, with later areas progressively sensitive to higher-level unpredictability – diverging from recent proposals, in both neuroscience and AI, that prediction operates primarily at higher levels of abstraction. At higher eccentricities, prediction effects are amplified but even V1 is tuned to high-level predictability, consistent with these prior accounts. Together, these results suggest that the visual system implements distinct prediction regimes across the visual field, thereby reconciling conflicting accounts of what visual cortex predicts.

---

Despite its apparent simplicity, seeing is among the brain’s most intricate computational tasks, requiring the transformation of noisy retinal signals into coherent percepts. A prominent idea in neuroscience and artificial intelligence (AI) is that prediction is fundamental to this process. For instance, theories of predictive processing propose that the brain constantly generates predictions about incoming sensory signals, with the discrepancy between predicted and actual input driving both perceptual inference and learning [1–3]. Prediction is also a key mechanism in models of self-supervised learning in (Neuro)AI, that describe different ways in which prediction of visual input can drive learning [4–9]. Yet despite its theoretical ubiquity, some fundamental questions about prediction in biological vision remain unresolved.

A first question concerns the automaticity of prediction. Both generative approaches to perceptual inference and models of self-supervised learning generally imply that prediction is continuous and ongoing — an inherent part of perceptual processing, or a key operation driving continual learning. By contrast, alternative frameworks cast prediction as a costly, strategic operation that is recruited only in specific situations: during particular tasks, or under challenging conditions when rapid feedforward processing proves insufficient [10–13]. A difficulty in arbitrating between these possibilities is that empirical evidence remains largely confined to artificial, prediction-encouraging experiments, such as cueing or oddball paradigms, with typically only a handful of possible stimuli and experimentally imposed predictions [14–16, 3, 17, 18]. While such studies demonstrate that sensory cortex *can* predict, they leave unclear whether these effects generalise to naturalistic stimuli, and whether prediction occurs automatically during natural perception.

A second question concerns the representational content of predictions: at what level of abstraction does the visual system predict – low-level features such as edges, higher-level features such as textures and objects, or a hierarchy spanning both? Classical predictive coding proposes hierarchical correspondence: early visual cortex is most sensitive to low-level unpredictability, with later areas progressively sensitive to higher-level unpredictability [1, 21, 22]. Reverse hierarchy theories imply the opposite: early areas resolve high-level unpredictability, while later areas are principally sensitive to residual low-level mismatch [11]. Finally, the brain may predict specifically at a single, high level of abstraction – a view supported by recent insights from self-supervised learning models in AI, where effective representations emerge from predicting more abstract features rather than redundant low-level details [7, 8, 23, 9, 24], and by an emerging empirical consensus suggesting that visual cortex is most sensitive to higher-level unpredictability [25– 29].

Advances in generative AI offer a way to address both questions. By using models that predict visual content from spatial context, one can estimate the inherent predictability of any visual stimulus at multiple levels of abstraction, thereby allowing to study sensory prediction during natural perception, without experimentally imposing predictions on the observer [25, 29–31].

Here, we combine high-resolution 7T fMRI recordings of human visual cortex responding to 73,000 natural images [20] with deep generative models that quantify spatial predictability at multiple levels of abstraction. We find that responses throughout V1–V4 are modulated by spatial predictability during natural scene viewing. In central vision, we observe a hierarchy of predictions that parallels the feature-encoding gradient [32], which is in line with classical predictive coding [1] but diverges from recent proposals that visual cortex predicts primarily at high levels of abstraction [25, 27, 26, 29]. At higher eccentricities, however, even early visual cortex is tuned to high-level predictability, consistent with these prior accounts. Together, these results suggest that the visual system implements distinct prediction regimes across the visual field, reconciling conflicting findings and constraining models of both predictive processing and self-supervised learning.

## Results

We examine the Natural Scenes Dataset (NSD; [20]), which comprises high-quality 7T fMRI recordings of 8 participants viewing a comprehensive set of 73.000 natural scene images, plus a battery of controlled stimuli to characterise tuning, including population receptive field (pRF) mapping. To probe sensory predictions, we used a deep generative model trained to predict image patches based on their surroundings, and then compare the predicted (reconstructed) image patch with the actually presented image patch to compute its spatial predictability (see Fig. 1; [25, 29]). For our main analyses, we focus on a single central patch of every image, and the voxels with pRFs within those central 2 ° of the visual field (see *Methods*). This results in a single unpredictability score per image, which we then compare to the brain response, for every image, for every selected voxel, across the visual hierarchy (V1-V4).

**Figure 1:**
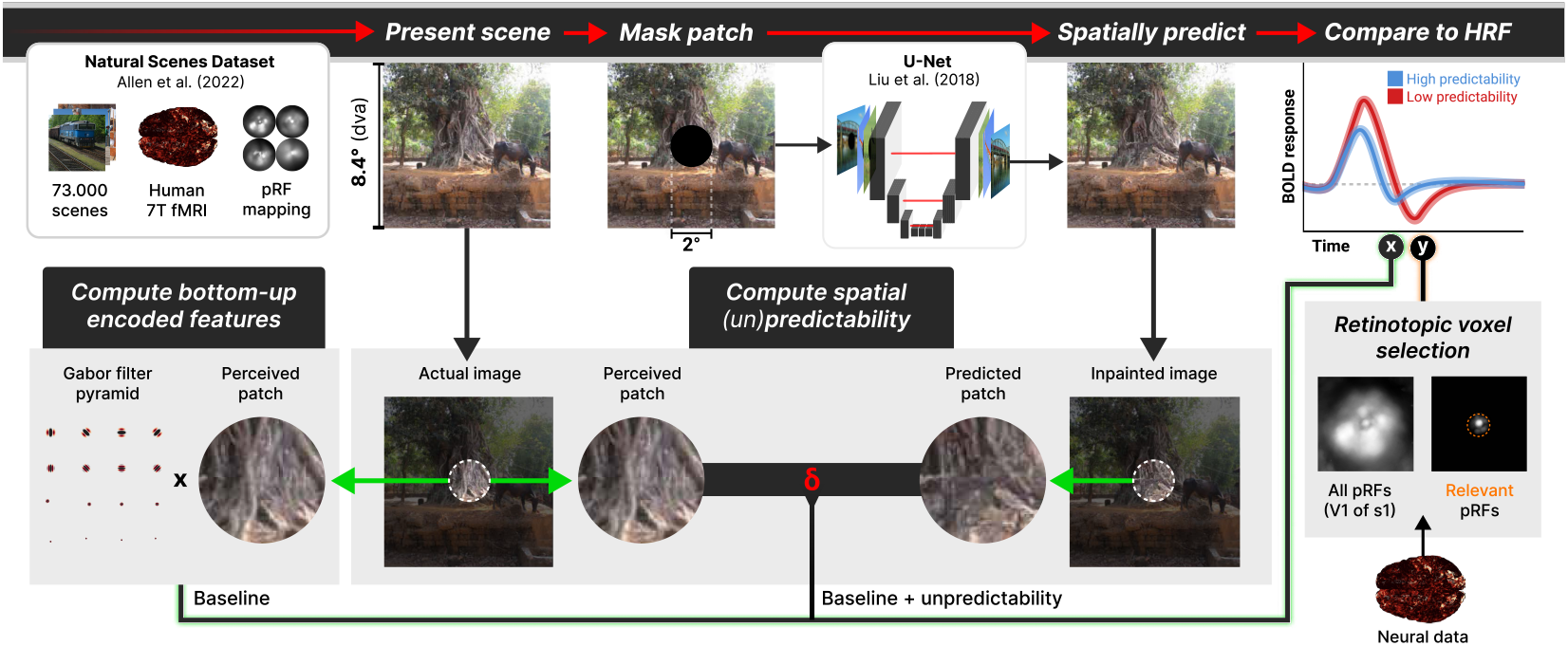
Empirical framework: data and modelling resources (white blocks) and analytical methods (grey blocks). We quantified the spatial predictability of natural image patches (1° visual angle radius) by comparing actual visual content with model-predicted reconstructions [19]. These predictability estimates were regressed on retinotopically (pRF) relevant fMRI BOLD responses from humans viewing 73,000 natural images (NSD [20]), controlling for low-level visual features (Gabor pyramid filter outputs).

### Spatial predictability modulates neural responses during natural vision

We first assessed whether visual cortex was indeed sensitive to spatial predictability during natural scene perception. We hypothesized that, if the brain generates spatial predictions, neural activity should reflect input predictability, with less predictable inputs evoking stronger responses.

To test whether spatial predictability indeed modulates neural responses, we first asked whether unpredictability explained unique variance, over and above a strong baseline model, which compiled a variety of bottom-up encoded local contrast features at different spatial scales, formalizing the hypothesis that natural vision does not involve spatial prediction. When we compared this baseline model to a regression model that additionally included each image’s unpredictability score, we found that we could consistently capture the brain response better, revealing additional explained neural variance (Fig. 2B; bootstrapped mean % Pearson’s *r* change versus baseline [95% CI], across subjects – in V1: 8.97% [6.97, 11.24]; V2: 7.53% [5.79, 9.45]; V3: 7.28% [5.84, 8.81]; V4: 7.38% [6.36, 8.68]), over and above the already substantial amount captured by the baseline model (Fig. 2C; bootstrapped mean Δ*r* [95% CI] versus shuffled control, across subjects – in V1: Δ*r* = 0.102 [0.088, 0.114], Cohen’s *d* = 8.94; V2: Δ*r* = 0.083 [0.076, 0.090], Cohen’s *d* = 14.53; V3: Δ*r* = 0.075 [0.067, 0.083], Cohen’s *d* = 9.11; V4: Δ*r* = 0.065 [0.055, 0.078], Cohen’s *d* = 7.01).

**Figure 2:**
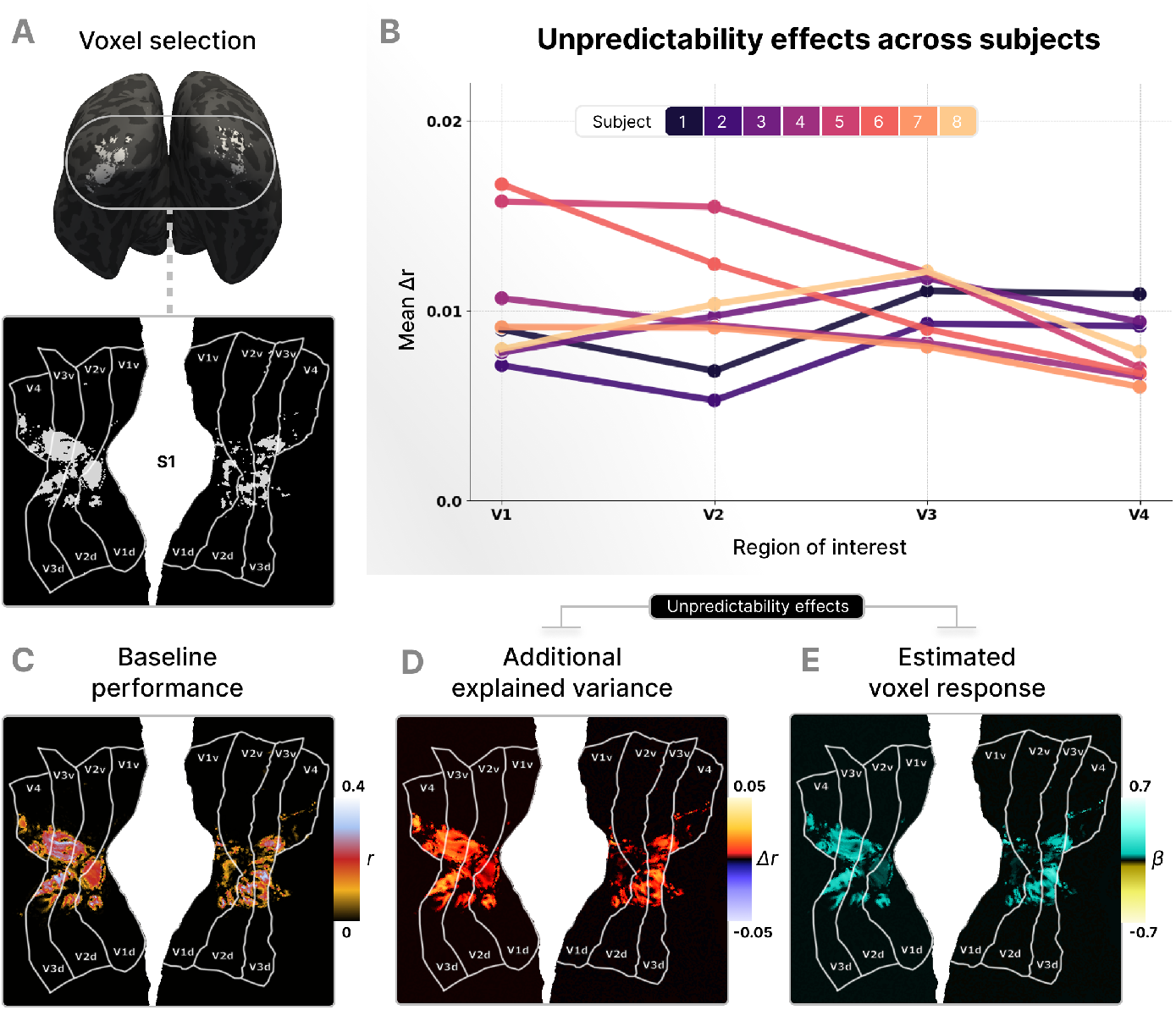
Spatial predictability modulates neural responses. A) Voxel selection of neural population with receptive fields inside the central image patch, based on pRF retinotopy (*subject 1*). B) Overall unpredictability effects. The additional explained neural response variance by spatial unpredictability features over and above the local-contrast baseline, indicated by the difference in cross-validated correlation coefficients for all subjects. C) Predictive performance of baseline model across the visual cortical surface indicates that local-contrast features account for substantial variance in BOLD activity of selected voxels. D) Additional explained variance reflects a consistent increase in model fit over and above baseline performance across the cortical surface. E) Direction of unpredictability effects aligns with expectation suppression, indicated by a consistent positive modulation of evoked neural response by unpredictability. Response modulation is expressed as the mean difference in estimated change of normalised neural activity for a 1*σ* increase of spatial unpredictability. All effects are consistent across subjects (SI appendix, Fig. S2), surface plots display data from a single subject representative.

This effect was consistent throughout the visual cortex and observed in each participant (Fig. 2B). Furthermore, we investigated the nature of this unpredictability effect, by examining the coefficients, revealing a positive modulation profile in line with expectation suppression: responses are stronger when sensory input is less predictable (Fig. 2E; bootstrapped mean cross-validated *β* coefficient [95% CI], across subjects – in V1: *β* = 0.049 [0.038, 0.063]; V2: *β* = 0.048 [0.041, 0.056]; V3: *β* = 0.048 [0.042, 0.054]; V4: *β* = 0.039 [0.034, 0.045]).

Together, these results indicate that the brain is sensitive to the spatial predictability of visual input during naturalistic perception. This suggests that visual processing involves the generation of context-based predictions about incoming visual input.

### A hierarchy of spatial predictions across the cortex

Thus far, our results demonstrated that human visual cortex is sensitive to an overall metric of spatial predictability. However, this does not address the representational granularity of the predictions: is visual cortex more sensitive to the predictability of lower-level or higher-level features – and how does this *unpredictability tuning* evolve across the visual system?

We contrasted three hypotheses on the tuning of sensory predictability across visual cortex (Fig. 3)C. First, following canonical predictive processing models [1, 2, 21] predictability tuning may mirror bottom-up feature tuning, with early visual areas sensitive to low-level unpredictability and later visual areas to higher-level unpredictability. Second, predictability tuning could exhibit a reverse effect, following reverse hierarchy theories [34, 35]. Finally, unpredictability sensitivity might be non-hierarchical, with sensory cortex being primarily sensitive to high-level unpredictability, across cortex, as suggested by recent findings [26–29, 25].

**Figure 3:**
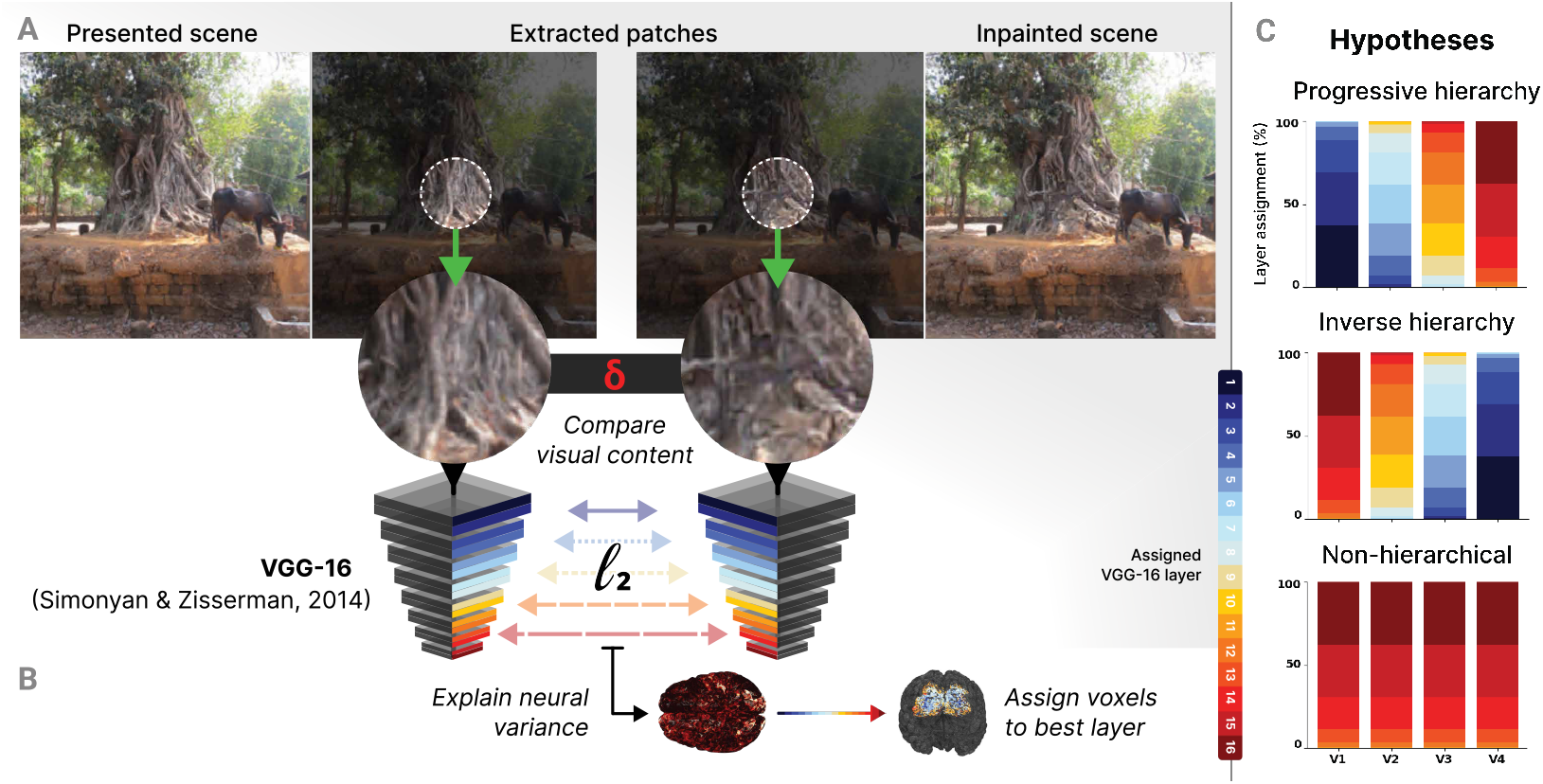
Spatial predictability at multiple levels of abstraction. A) Perceived and model-inpainted natural scene patches were submitted to a deep vision-trained neural network (VGG-16) [33] and compared by computing the Euclidean distance (*l*_2_) between patches’ increasingly abstract feature representations at each convolutional and dense layer of the network. B) Cross-validated ridge regression of visual cortical responses on spatial unpredictability. Voxels were assigned to the level of abstraction (i.e. VGG-16 layer) at which spatial unpredictability features explained most neural variance above and beyond the local-contrast baseline. C) Hypothesised spatial unpredictability tuning profiles.

Before establishing unpredictability tuning, we first assessed the feature tuning, by constructing voxel-wise encoding models based on a pre-trained Convolutional Neural Network (CNN), and establishing for each voxel which layer of the CNN best predicted that voxel (layer assignment, [32]). The results (SI Appendix, Fig. S3) recapitulated the classical tuning hierarchy, where early visual cortex is best accounted for by early CNN layers (representing low-level features) and higher-level visual areas become progressively sensitive to more abstract features [36], with 41% of V1, 60% of V2, 75% of V3, and 87% of V4 most sensitive to high abstract (> conv layer 8) features.

Having derived this visual cortical tuning reference for bottom-up features, we proceeded by analyzing its tuning profile for spatial predictability. To this end, we computed spatial unpredictability at multiple levels of feature abstraction (i.e. VGG-16 feature space) (Fig. 3A) and assessed their individual neural predictivity (Fig. 3B). This allowed us to ask, for every voxel across the visual hierarchy, analogously to the feature-encoding analysis, what level of unpredictability it was most sensitive to.

Strikingly, this revealed a consistent hierarchy of predictability sensitivity (Fig. 4) across the visual system (SI Appendix, Fig. S4), in which higher-level visual areas become progressively sensitive to more abstract levels of spatial predictability, with 23% of V1, 32% of V2, 45% of V3, and 73% of V4 most sensitive to high abstract (> conv layer 8) predictability. This tuning gradient for sensory predictability parallels the established bottom-up feature encoding hierarchy and aligns with hierarchical accounts of predictive coding in the cortex, which postulate a hierarchy of predictions, in which every cortical area sends top-down predictions about the expected features encoded at the level below [37, 2, 31, 38, 39, 3, 40]. Interestingly, however, this hierarchical gradient diverges critically from recent observations suggesting that visual cortex – even early areas –predominantly predicts at higher levels of abstraction [25, 29, 27, 28].

**Figure 4:**
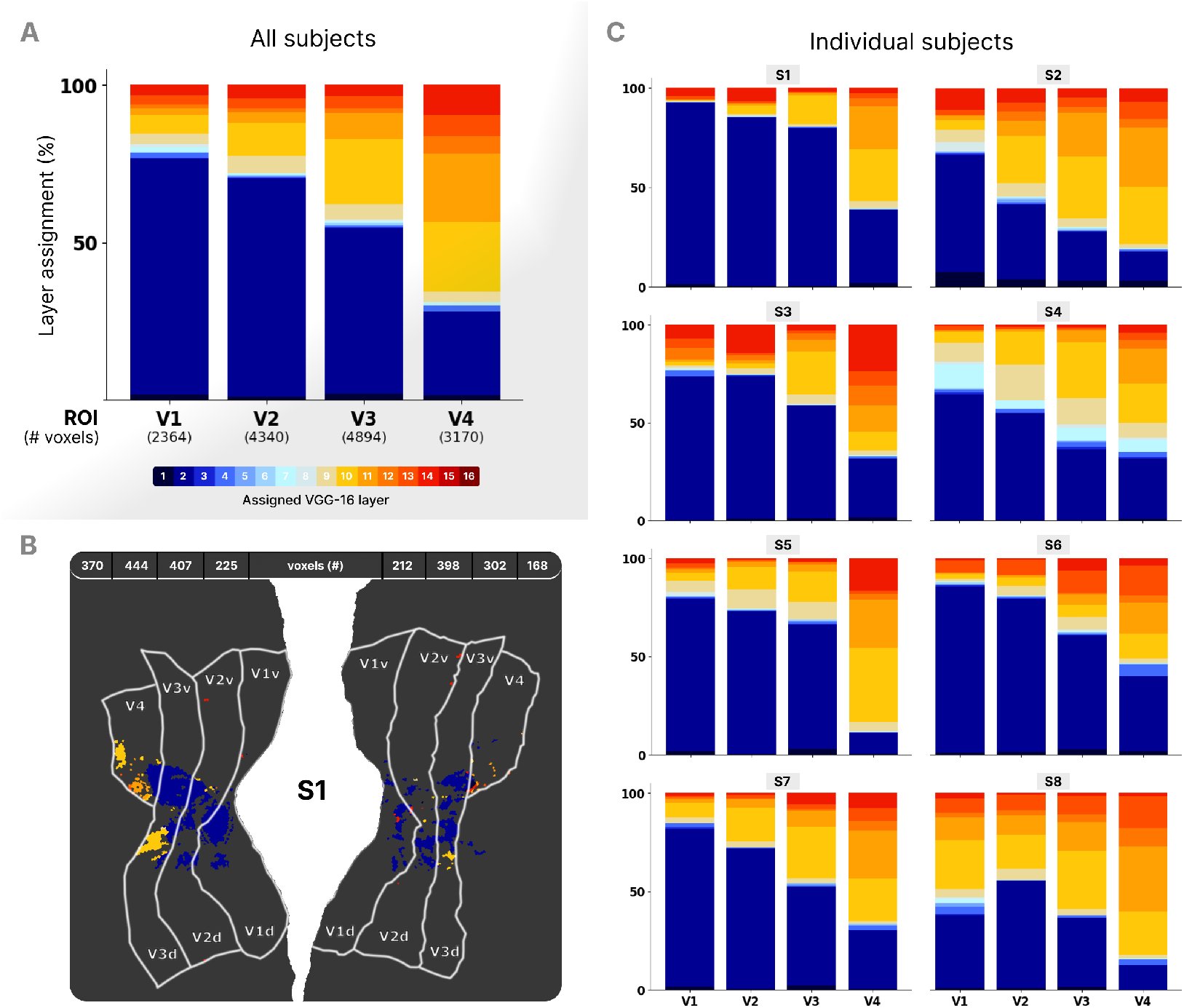
Visual cortical sensitivity to distinct levels of spatial unpredictability. A) Voxel assignment to the abstract level at which spatial predictability explains most neural variance over and above baseline features, aggregated over subjects. B) Spatial predictability sensitivity across the flattened visual cortical surface, subject-specific. C) Robustness of spatial prediction tuning profile displayed by single-subject voxel assignments.

### Eccentricity modulates both strength and hierarchy of spatial prediction

What might account for this divergence? One notable difference is that the most directly comparable prior studies examined neural populations with more peripheral receptive fields: mouse visual cortex, whose RFs span 20–30° [29], and macaque V1 at ±5° eccentricity [25]. Because whole-brain fMRI provides coverage across the visual field, we could test whether prediction differs as a function of RF eccentricity, and hence sensory reliability.

This allowed us to test two hypotheses. First, following accounts proposing that reliance on prediction scales with sensory uncertainty [2, 41, 42], prediction effects should be stronger at higher eccentricities, where sensory resolution is lower. Second, if the divergence between our results and prior work reflects eccentricity-related RF size, parafoveal voxels should behave more like the prior reports – that is, showing sensitivity to high-level unpredictability even in early visual areas.

To test this, we repeated the analysis at three para-foveal patches (2° eccentricity) and compared effects to the foveal patch, within each participant. As predicted, spatial predictability effects were systematically stronger in parafoveal compared to foveal voxels (Fig. 5E). This pattern is found in all participants (SI Appendix, Fig. S5) and every visual cortical area (bootstrapped mean % Pearson’s *r* change versus baseline [95% CI], across subjects – in V1: fovea 8.97% [6.97, 11.24], parafovea 65.60% [58.95, 73.14], difference: +56.64%, Cohen’s *d* = 7.86; V2: fovea 7.53% [5.79, 9.45], parafovea 55.39% [51.69, 59.73], difference: +47.86%, Cohen’s *d* = 11.02; V3: fovea 7.28% [5.84, 8.81], parafovea 51.86% [43.64, 60.58], difference: +44.58%, Cohen’s *d* = 5.44; V4: fovea 7.38% [6.36, 8.68], parafovea 55.55% [44.13, 69.21], difference: +47.72%, Cohen’s *d* = 3.54).

**Figure 5:**
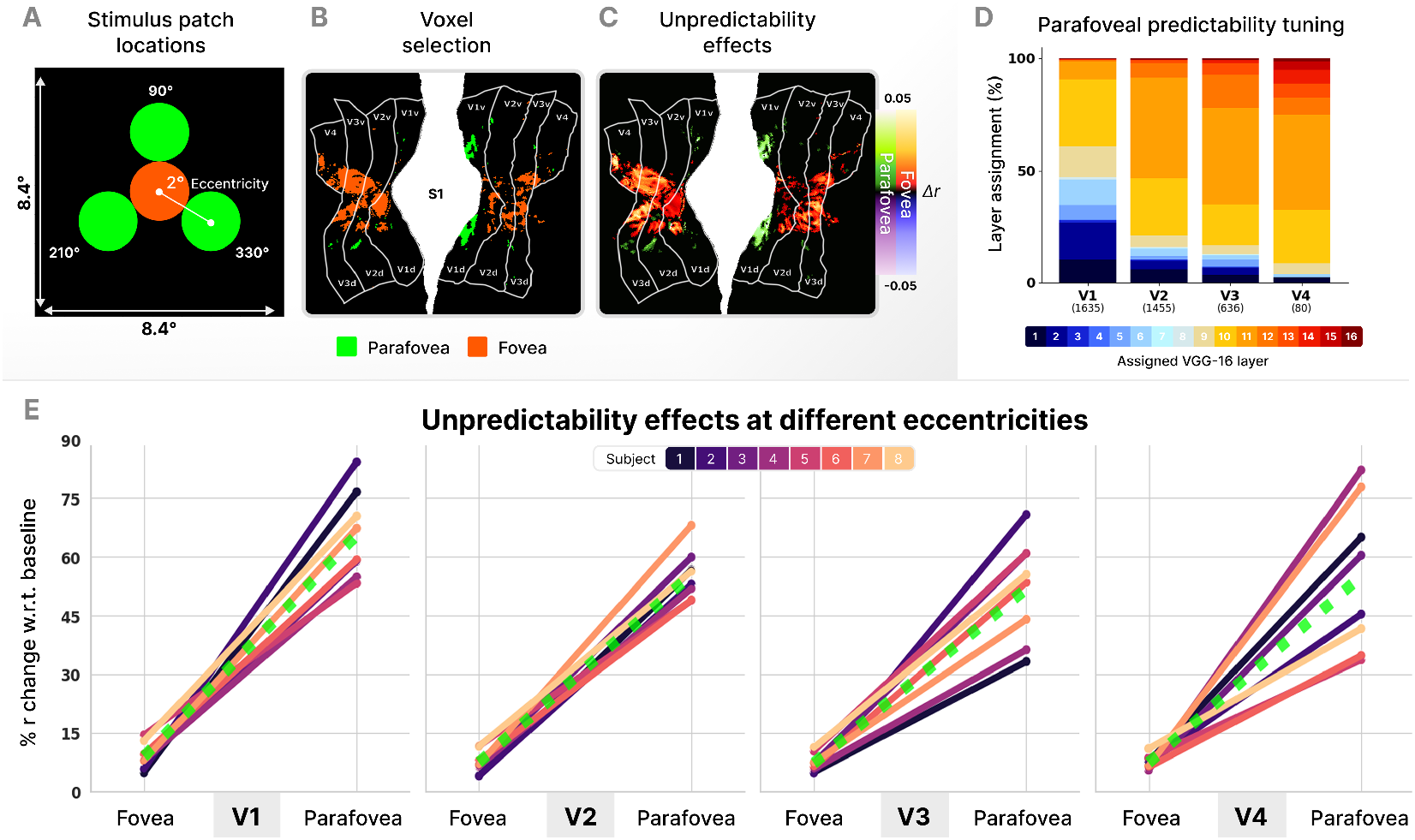
Sensitivity to spatial predictability is stronger when sensory reliability is lower. A) Relative location of analysed natural scene patches. Parafoveal analyses considered aggregated data from a triplet of equally sized parafoveal patches with fixed eccentricity and complementary angles. B) Selection of voxels with sufficiently similar pRFs to a parafoveal patch, displayed across the visual cortical surface of a single subject. C) General spatial predictability effects in the (para)fovea, expressed as uniquely explained neural variance over and above baseline features. D) Voxel assignment to the abstract level at which spatial predictability uniquely explains most neural variance beyond the baseline. E) Sensitivity to spatial predictability is systematically stronger in the parafovea than in the fovea, expressed as percentage change with respect to neural predictivity of the baseline model (SI Appendix, Fig. S6).

We then examined the predictability tuning in para-foveal voxels. While we still observed a gradient – with higher visual areas sensitive to higher-level predictability (Fig. 5D; high abstract [> conv layer 8] predictability tuning: V1 53%; V2 85%; V3 87%; V4 96%) – there was a marked shift towards high-level predictability compared to central vision. Even V1 was most sensitive to high-level predictability in the parafovea, in contrast to its low-level tuning in the fovea. Critically, this shift was specific to predictability: sensitivity to image features showed no such eccentricity-dependent change (SI Appendix, Fig. S7, bottom).

Together, these results indicate that the hierarchical prediction gradient we observe may be specific to high-acuity central vision. At higher eccentricities, prediction operates predominantly at higher levels of abstraction, consistent with prior animal work, and potentially reflecting distinct functional regimes of spatial prediction across the visual field (see *Discussion*).

## Discussion

We quantified the spatial predictability of image patches across tens of thousands of natural images and related these to high-resolution fMRI responses throughout human visual cortex. This revealed robust spatial prediction effects across the visual hierarchy, with more predictable patches evoking weaker responses. In central vision, we observe a hierarchy of predictions: early visual areas are most sensitive to low-level unpredictability, while later areas become progressively sensitive to higher-level unpredictability; a gradient that parallels bottom-up feature encoding. At higher eccentricities, where sensory input is less reliable, we find that predictability effects are stronger – but we do not observe the same hierarchy of predictions as in central vision. Instead, we observe a sensitivity for more high-level predictability, even in V1. Together, these results underscore the ubiquity of prediction across cortex, and suggest that visual cortex implements distinct regimes of prediction across the visual field – with implications for theories of predictive processing and self-supervised learning.

While prediction in neuroscience is often studied using temporal predictions [16, 43], we focus on spatial prediction, which is the kind of prediction described by canonical predictive coding models [1, 21], and connects our findings to a rich history of work on spatial context effects in vision. The suppressive effect we observe is conceptually most similar to surround suppression, where responses to stimuli in the receptive field are reduced by coherent content in the surround [44, 45] – a phenomenon commonly explained as reflecting redundancy reduction or predictive coding [46, 47, 1]. However, the opposite (surround enhancement) is also also well documented [45] and may be more prevalent for natural stimuli [48]. Methodological differences further complicate direct comparisons, as the classical analysis is based on matching RF content to surround content, which is not what we do. Conceptually, though, the suppressive effect of predictability we observe is in line with predictive or efficient coding accounts of surround modulation – but departs from classical surround suppression in two respects. First, we find robust predictability effects in the central 2°, where surround suppression is typically weak or absent [49, 50]. Second, at higher-eccentricities, we observe suppression strongest to higher-level predictability even in areas tuned to low-level features – whereas classical surround suppression is strongest when the surround matches the neuron’s preferred features [44, 51].

A distinct influential line of work has studied spatial prediction using occlusion: analysing neural activity in neurons or voxels responding to occluded part of a visual scene, and observing that these ‘blind’ regions nonetheless encode information about the occluded scene content [52–56]. While this approach isolates feedback by removing feedforward input, ours provides a complementary view, by showing that prediction also operates when bottom-up input is present, and allowing to model the content of predictions across the visual field.

Our finding of a hierarchy of predictions in foveal vision appears to conflict with an emerging consensus suggesting that visual cortex – even in early areas – predominantly predicts at higher levels of abstraction [28, 25, 26, 57, 29]. However, as our eccentricity analysis revealed, this apparent contradiction might reflect different regimes of prediction under different conditions of reliability. The spatial prediction studies most comparable to ours [25, 29] examined neurons with RF properties most similar to human (near)-peripheral receptive fields, and our parafoveal analysis recapitulated their findings: a shift towards high-level predictability tuning, with no corresponding change in feature tuning. A similar logic may apply to studies using temporal prediction, which also showed high-level primacy [28, 26, 57]. These involve associations between discrete stimuli learned specifically for the experiment, which induce predictions that are inherently coarser than predictions about fine-grained spatial structure based on extensive experience with the world (as we study here). Together, this suggests a unifying principle: under uncertainty, visual cortex defaults to high-level prediction – but under high reliability (of which spatial prediction in the fovea is an extreme case), predictions can extend all the way down to low-level features.

Why would the brain be engaged in such different types of prediction simultaneously? One possibility is that they reflect distinct functional demands. Prediction can serve at least two purposes: inference and learning. For inference (recognition) predicting at multiple levels of abstraction is computationally useful, since ambiguities occur at multiple levels (e.g. when a faint edge is recognised by virtue of being part of a critical object boundary; [21, 13]). For learning, by contrast, prediction at a single level may suffice. Indeed, advances in self-supervised learning have shown that in vision, predicting high-level features drives more effective representation learning [7, 8, 58, 24, 23]. This functional asymmetry maps naturally onto the visual field. Foveal vision is uniquely suited to detailed recognition and discrimination – tasks requiring multi-level inferential precision [59] – while peripheral vision, with its coarser resolution, primarily supports gaze guidance and rapid scene understanding [11]. This focus on inference may explain why hierarchical prediction might be specific to central vision; for peripheral processing, high-level prediction alone may be both sufficient and efficient.

Together, our results highlight the ubiquity of prediction in human vision and reveal a hierarchy of predictions across visual cortex that parallels the hierarchy of feature encoding, but appears to be specific to central vision. This suggests that the answer to “what does visual cortex predict?” may depend on where in the visual field one looks – and that even the most general principles of computation must ultimately come into focus in the fovea.

## Materials and methods

### Stimuli and dataset

We analysed the stimuli and neural data from the Natural Scenes Dataset (NSD; [20]). This open access dataset compiles whole-brain 7T BOLD fMRI recordings from 8 human subjects perceiving tens of thousands natural scenes in a naturalistic viewing paradigm. These images were presented for 3 s and spanned a range of 8.4°x 8.4°visual angle. Subjects performed a continuous recognition memory task in which they reported whether the current image had been shown before. We used the neural data from the main experiment for our main analysis and data from the population receptive field (pRF) experiment for retinotopy-based ROI definition. More specifically, we used the single trial beta coefficients – estimates of the haemodynamic response function (HRF) amplitude – from GLMsingle [60] as a proxy for neural activity. We analysed all of the natural scene stimuli, except for a subset held out for cross-validation of regression models. All reported effects reflect held-out model performance.

### Population receptive field mapping, voxel selection

To enable a single unpredictability estimate per image to be compared against neural responses, we restricted our main analyses to a central patch (1° radius) of each stimulus image, and selected only voxels whose receptive fields fell within this patch. Population receptive fields (pRFs; [61]) were estimated using compressive spatial summation (CSS) models [62] from the NSD’s dedicated pRF mapping experiment, and visual cortical areas V1–V4 were defined based on the retinotopic parcellation provided with NSD. We selected voxels with a pRF radius between 0.15° and 1.0° visual angle, and a pRF centre within (1− pRF radius)° of the patch centre, ensuring that each selected voxel’s receptive field fell entirely within the analysed patch. This guarantees that the neural responses entering our analyses are specifically attributable to the image content for which we compute features and predictability.

### Spatial predictability inpainting model

To compute spatial predictability, we used a partial convolution U-Net (PConvUNet; [19]) that reconstructs (or inpaints) the masked-out image patches from their surroundings. It was pre-trained on ImageNet with non-contiguous irregular holes as in [19], and fine-tuned trained on the Places2 dataset [63] for 20,000 iterations using contiguous circular masks to match our patches of interest (architecture and training details in SI Appendix, section S1 and S2). For all 73,000 NSD images, we erased each patch with a boolean mask and had it painted back in from what remained. The resultant reconstructions constituted the model’s spatial prediction: the most probable patch content provided its visual context.

### Spatial predictability analysis

To estimate patch unpredictability, we compared these reconstructions (spatial predictions) with actual patches, for which we encased each patch in a square 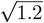 times its diameter and computed the Euclidean distance (*l*_2_) between their increasingly abstract feature representations at each convolutional and dense layer of VGG-16, a deep vision-trained neural network with representational resemblance to the human ventral visual stream [32, 64]. We examine the visual difference between patches in the latent space of a CNN because it robustly captures perceptually relevant features at distinct granular levels, regardless of pixel-level trivialities [65].

To test whether visual cortex is sensitive to spatial predictability in general (Fig. 2), we aggregated these levels for an overall metric of spatial predictability by simply taking the mean of all levels’ perceptual distances. For the more fine-grained analyses of predictability tuning (Fig. 4), we used the layer-specific perceptual distances to test for relative effects of predictability at different abstract levels. As these levels follow the sequential processing layers in VGG-16, nearby predictability estimates are more correlated than estimates at levels farther apart. Importantly, besides such similarity in adjacent layers, the dissimilarity of spatial predictability is also clearly structured throughout. Estimates’ layer-wise divergence follows a systematic and interpretable pattern, most apparent in textured images (SI Appendix, Fig. S1), and in line with seminal work on the mathematical dissociation between low and high-level visual features [66].

### Baseline model

To isolate contributions of spatial predictability above and beyond feedforward visual drive, we constructed a Gabor filter pyramid baseline [67] — a comprehensive model of local contrast features encoding orientation (0°, 45°, 90°, 135°) andspatial frequency (0.5, 1, 2, 4 cycles/dva) across the patch region. Filter outputs were projected onto their top 100 principal components (∼93% variance), together constituting the baseline feature representation. This baseline formalises the null hypothesis that any apparent, isolated spatial predictability effects can be explained by local image statistics rather than context-based visual predictability.

### Regression

We used ridge regression with standard 5-fold K-fold cross-validation to relate derived image features and predictability to voxel-wise HRF beta coefficients, quantifying performance as the cross-validated Pearson correlation (*r*) between actual and predicted neural responses across folds. Folds were contiguous trial splits in presentation order to prevent temporal leakage. We compared a baseline model of local contrast features (see *Baseline Model*) to an unpredictability model appending a single spatial unpredictability score. The difference Δ*r* (r_unpre-dictability −r_baseline) indexed the unique variance explained by predictability, over and above the baseline. The regression coefficient *β* of the unpredictability scores estimates the response change per 1 *σ* increase in unpredictability; a positive *β* is consistent with expectation suppression. For predictability tuning, we fitted separate models for each VGG-16 layer’s unpredictability score and identified the layer maximising Δ*r* per voxel, enabling a relative comparison with the bottom-up feature encoding hierarchy (see *Bottom-up feature encoding reference*; SI Appendix, Fig. S3).

### Bottom-up feature encoding reference

To acquire a voxel tuning reference for bottom-up encoded features, we assessed to what abstract level of visual features voxels in visual cortex are most sensitive. For this, we extracted VGG-16 feature representations of all natural scenes from its convolutional and dense layers, and reduced their dimensionality to the 500 principal components best capturing their variance. We then examined for each voxel at which layer these feature representations had most predictive power for its activity, assigning them to the layer yielding the highest cross-validated Pearson’s *r*, compared to a shuffled model alias.

### Eccentricity analysis

To test whether predictability effects scale with sensory reliability, we repeated all analyses for three parafoveal patches (1° radius; 2° eccentricity; 90°, 210°, 330° polar angle). Voxel selection criteria were identical to the foveal analysis, including the size of our patch of interest – to ensure the same quality of reconstruction and resulting unpredictability features, which depends on the size of the area to reconstruct. While our patch area remained constant, receptive fields enlarge with eccentricity and along the visual cortical hierarchy. Resultingly, voxels gradually outgrew RF-size selection criteria in both ways, reducing their eligibility. Due to this, parafoveal analyses involve less voxels than foveal analyses, despite having triple the patches, most particularly in V4.

For every separate parafoveal patch, we computed spatial unpredictability and assessed how much neural variance it explained. The average of these neural predictivities served as aggregate metric for parafoveal effects, and compared within subjects to foveal effects. We assessed overall spatial unpredictability effects as well as unpredictability tuning (layer assignment). Importantly, we used the same inpainting model, patch sizes, and voxel selection criteria regardless of patch eccentricity. As such, we used the same procedures to acquire unpredictability estimates and neural populations across foveal and parafoveal analyses. Because of this, any shift in tuning reflects a change in the brain’s prediction regime rather than a change in stimulus statistics, or sample-specific receptive field properties.

## Data and code availability

The Natural Scenes Dataset (NSD) is openly available and can be accessed via https://naturalscenesdataset.org. The code-base of the current study is found at https://github.com/WiegerScheurer/prfpred-nsd.

## Acknowledgements

We thank Emily J. Allen and colleagues [20] for creating and publicly sharing the NSD. We also thank all the developers responsible for the open source software supporting our analyses.

## Author contributions

M.H. designed research; W.H.S. performed research; W.H.S. and M.H. contributed new reagents/analytic tools; W.H.S. analyzed data; and W.H.S. and M.H. wrote the paper.

## SI Appendix

### Supplementary methods

#### S1 Inpainting model architecture

The PConvUNet encoder consisted of eight blocks: the first one used a kernel of 7 × 7, the second and third used 5 × 5 kernels, and later blocks used 3 × 3 kernels. Each encoder block applied a partial convolution, batch normalisation (except for the first layer), and ReLU activation function. After each partial convolution, the binary validity mask was updated by marking all locations where at least one valid pixel was covered by the filter’s receptive field, progressively expanding the inpainted area as features propagated through the encoder. Feature responses were normalised by the ratio of valid (unmasked; informative) to total pixels within the kernel footprint, preventing regions from biasing feature statistics.

The decoder mirrored the encoder’s eight blocks, each of which performed nearest-neighbour upsampling, concatenation with the corresponding encoder feature map via a skip connection, partial convolution, batch normalisation, and LeakyReLU activation. Skip connections preserved spatial structure that would otherwise be lost through downsampling, reducing blurring and structural artefacts. Encoder weights were initialised from VGG-16 pretrained on ImageNet. During this training, encoder batch normalisation layers were frozen to preserve pretrained feature extraction while decoder parameters were free to be updated.

#### S2 Model training and loss function

The model was trained on the Places2 scene dataset [1] for 20,000 iterations (batch size 64) using the Adam optimiser (learning rate 5×10^−4^, *β*_1_ = 0.9, *β*_2_ = 0.999). Circular masks matched the size and shape of the image patches used in our analyses, ensuring the model was optimised specifically for receptive field-scale occlusions.

The training objective combined five distinctly weighted components:

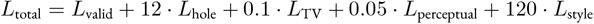

*L*_valid_ and *L*_hole_ are per-pixel ℓ_1_ reconstruction losses on unmasked and masked regions respectively. *L*_TV_ is a total variation penalty promoting spatial smoothness. The perceptual loss,

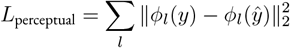

measures representational similarity across VGG-16 layers *ϕ*_*l*_, and the style loss,

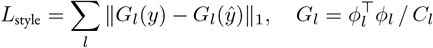

compares Gram matrices *G*_*l*_ capturing texture statistics, where *C*_*l*_ is the number of channels at layer *l, y* the ground-truth image, and *y*ŷthe reconstruction. The combined perceptual and style terms penalise representational discrepancies beyond pixel-level errors, reducing perceptual artefacts typically found in purely pixel-wise approaches. Full derivations follow Liu *et al*. [2].

#### S3 Cross-validation specifics

Standard 5-fold K-fold cross-validation was implemented using scikit-learn’s `KFold(n_splits=5, shuffle=False)`. Trials were partitioned contiguously in their natural presentation order across sessions. For each voxel, ridge regression models (α = 0.1) were trained on 4/5 of trials and evaluated on the held-out fold; Pearson correlations were computed between observed and predicted responses and averaged across folds.

#### S4 Model fitting procedure

Separate ridge regression models were fit voxel-wise for each ROI (V1–V4). Baseline models used the 100 leading principal components (PCs) of z-scored Gabor wavelet features (capturing ∼93% variance). Unpredictability models appended a single z-scored MSE reconstruction loss score per VGG-16 layer. Models were evaluated individually for each of the 13 convolutional + dense layers. The final unpredictability coefficient *β* was extracted from the corresponding regressor column and averaged across CV folds.

#### S5 Data preprocessing

Both X (features) and y (HRF beta coefficients) matrices were z-scored per voxel/feature prior to model fitting (fit_intercept=False) to center data and make *β* coefficients interpretable. Voxel selection was done using the VoxelSieve class, with pRF size constraints `(min_size=0.15, max_size=1, patchbound=1)`.

#### S6 Parafoveal effect metric

We compared spatial unpredictability effects in the parafovea and fovea within subjects based on a standardised metric (% change w.r.t. baseline performance) instead of Δ*r* for uniquely explained variance, because variability in signal to noise ratio across neural populations can artificially in-/ deflate relative differences in effect magnitude, which are derived from GLM-based estimates of the haemodynamic response. In other words, for this *inter*-areal comparison (fovea versus parafovea) of *intra*-areal (baseline versus baseline + unpredictability) effects, we control for potential *inter*-areal differences in effect scale by using a relative instead of an absolute difference metric.

#### S7 General technical details

Analyses were performed in Python 3.11 with scripts and IPython notebooks [3], supported by Shell interface scripts for workflow integration and coordination. Visualisation and transformation of neural data was performed using FreeSurfer [4], FSL [5], Pycortex [6], and a selection of publicly available NSD code repositories.

**Figure S1:**
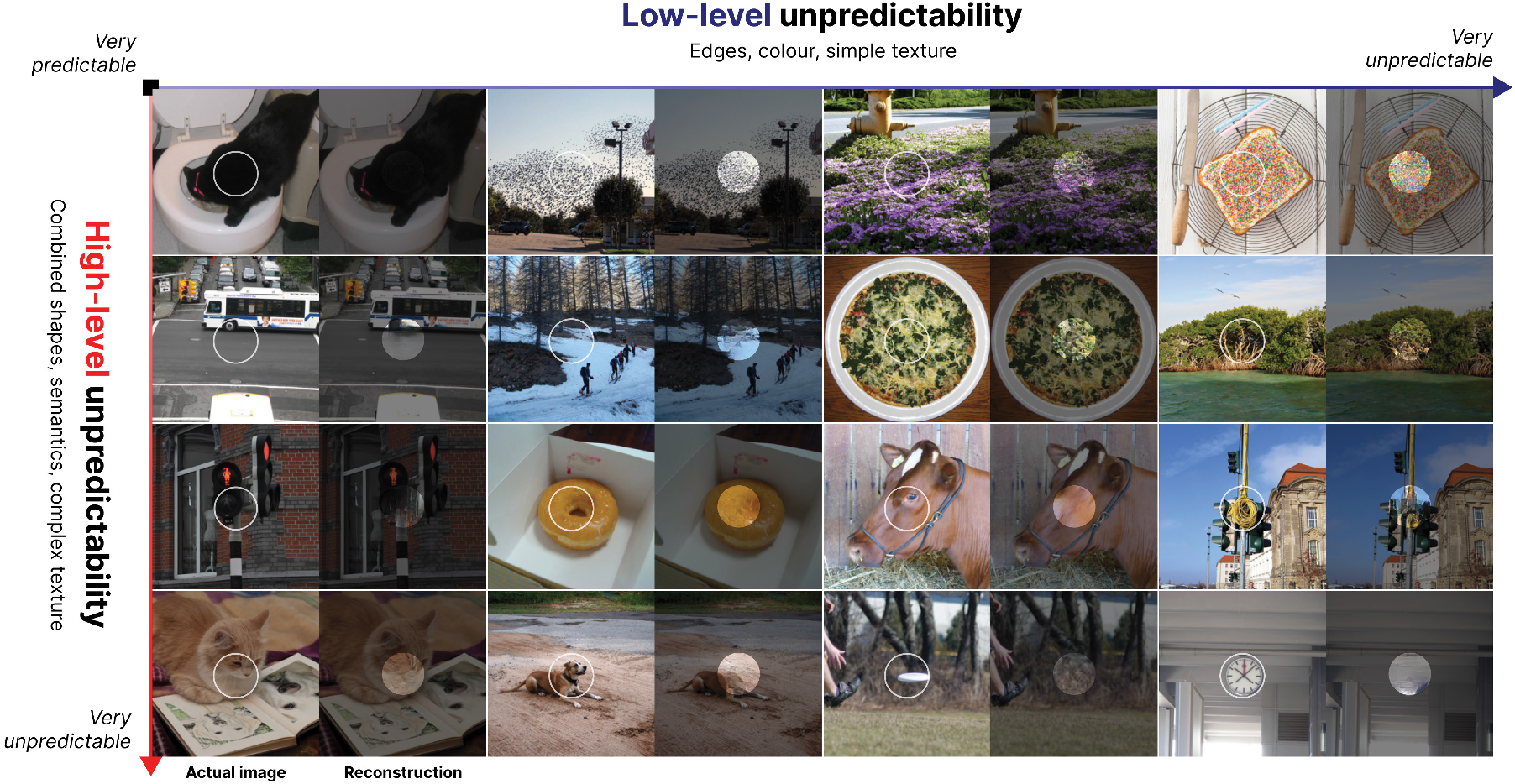
The distinction between spatial unpredictability at different levels of abstraction. Example stimuli that illustrate the distinction between spatial unpredictability at low versus high levels of abstraction. Every image pair reflects a distinct combination of low-level (horizontal axis) and high-level (vertical axis) spatial unpredictability — the difference in central patch content between the actual (left) and reconstructed (right) image. The extremes of the grid illustrate intuitive dissociations. Patches with low unpredictability on both axes typically contain simple, predictable structures like uniform regions or clear edges. Patches with high low-level but low high-level unpredictability are often natural textures: their fine-grained details are difficult to predict, but their overall structure (e.g., foliage, gravel) is easily inferred from context. Complex objects or object parts that stand out from the surround and are fully covered by the receptive field represent the opposite extreme, where both high-level and low-level features are highly unpredictable. The rarer case of low low-level but high high-level unpredictability can occur when edges or low-level features are predictable but nonetheless do not create a coherent high-level feature (e.g. object part).

**Figure S2:**
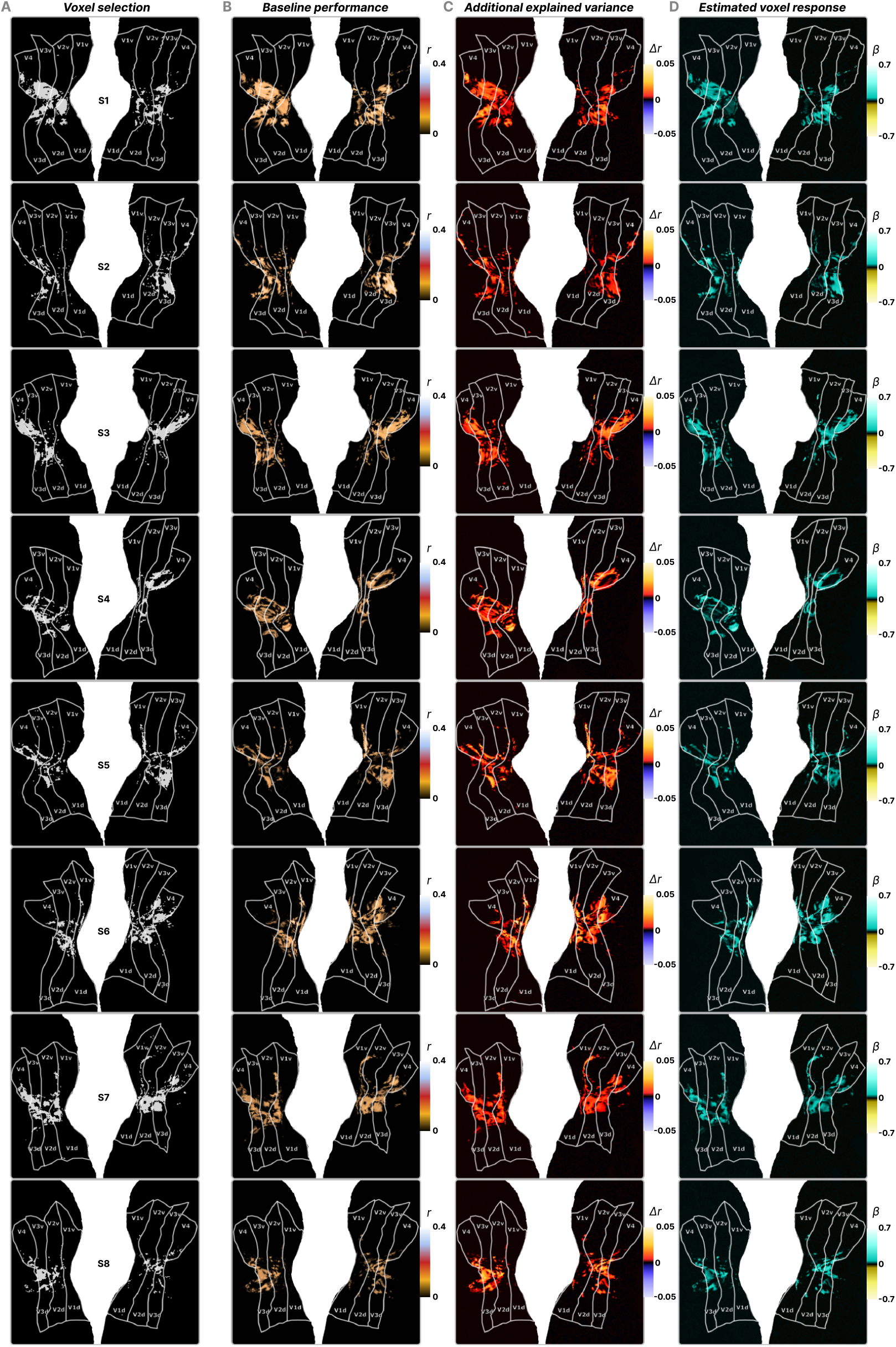
Consistency of local contrast and spatial unpredictability effects across the visual cortex in all subjects. A) Selection of voxels with pRFs sufficiently overlapping the foveal patch of interest. B) The baseline model consisting of local contrast features explains substantial neural variance in each individual subject, indicated by the Pearson’s correlation (*r*) between the actual and model-estimated amplitude of evoked responses. C) Overall metric of spatial unpredictability explains neural variance over and above the baseline model, evidenced by the difference in *r* (Δ*r*) between the baseline model and a variant with an added spatial unpredictability regressor. This is evident in every subject, where values are consistently positive across the full cortical surface that ROIs span. D) Neural responses become stronger when the visual content inside the patch is more unpredictable, indicated by the consistently positive *β* coefficients in the test sets from 5-fold cross-validated ridge regression models.

**Figure S3:**
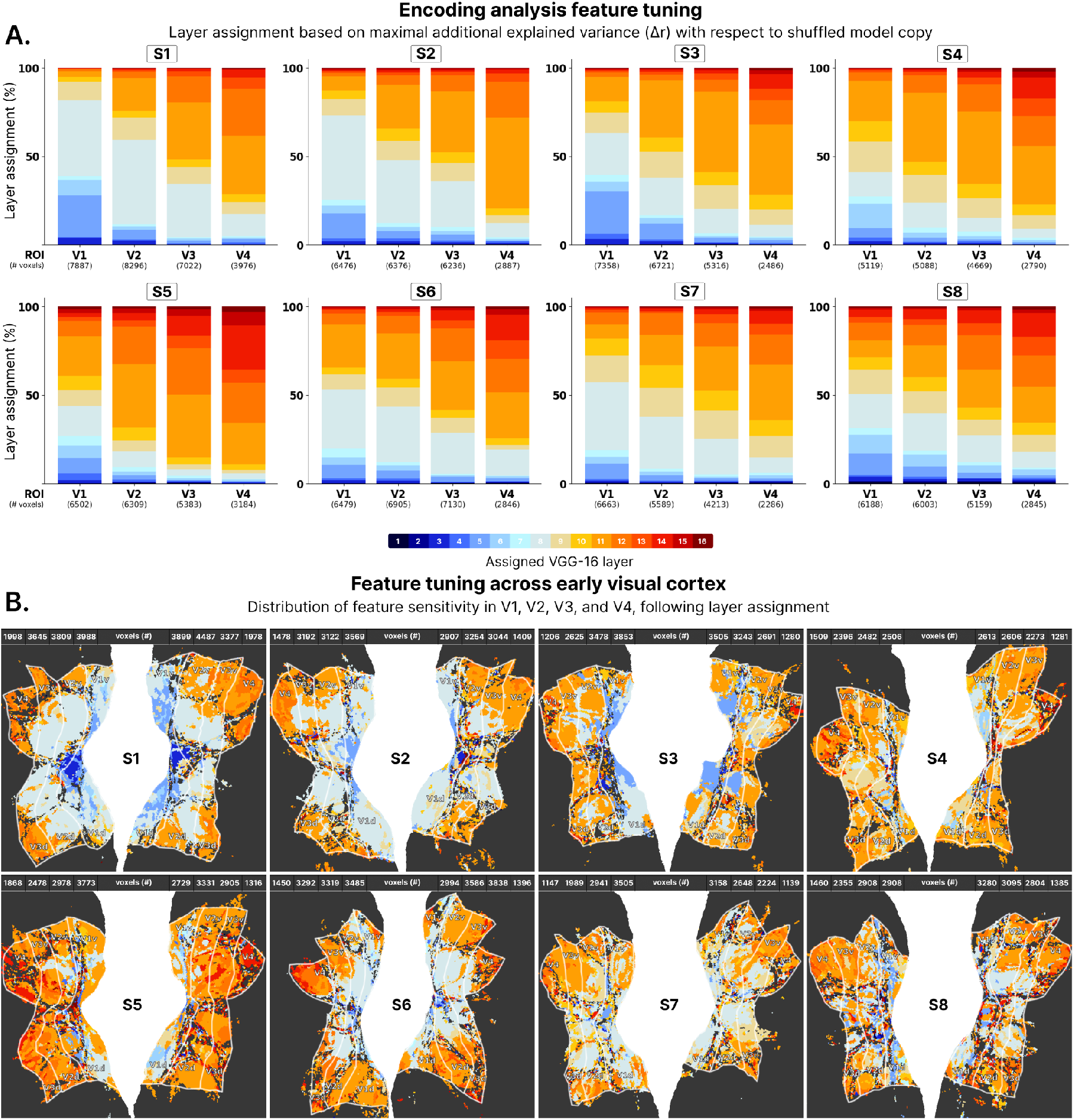
Visual cortex is hierarchically tuned for bottom-up encoded visual features. A) Voxel assignment to the abstract level at which VGG-16 feature representations of natural scenes (full view) predict neural variance best, per subject. In each individual, visual cortical areas are progressively tuned for more abstract visual features. B) Distribution of voxel assignment across flattened visual cortical surface, demonstrating the systematic hierarchical gradient of feature tuning along the cortex, with later areas progressively tuned for more abstract visual features.

**Figure S4:**
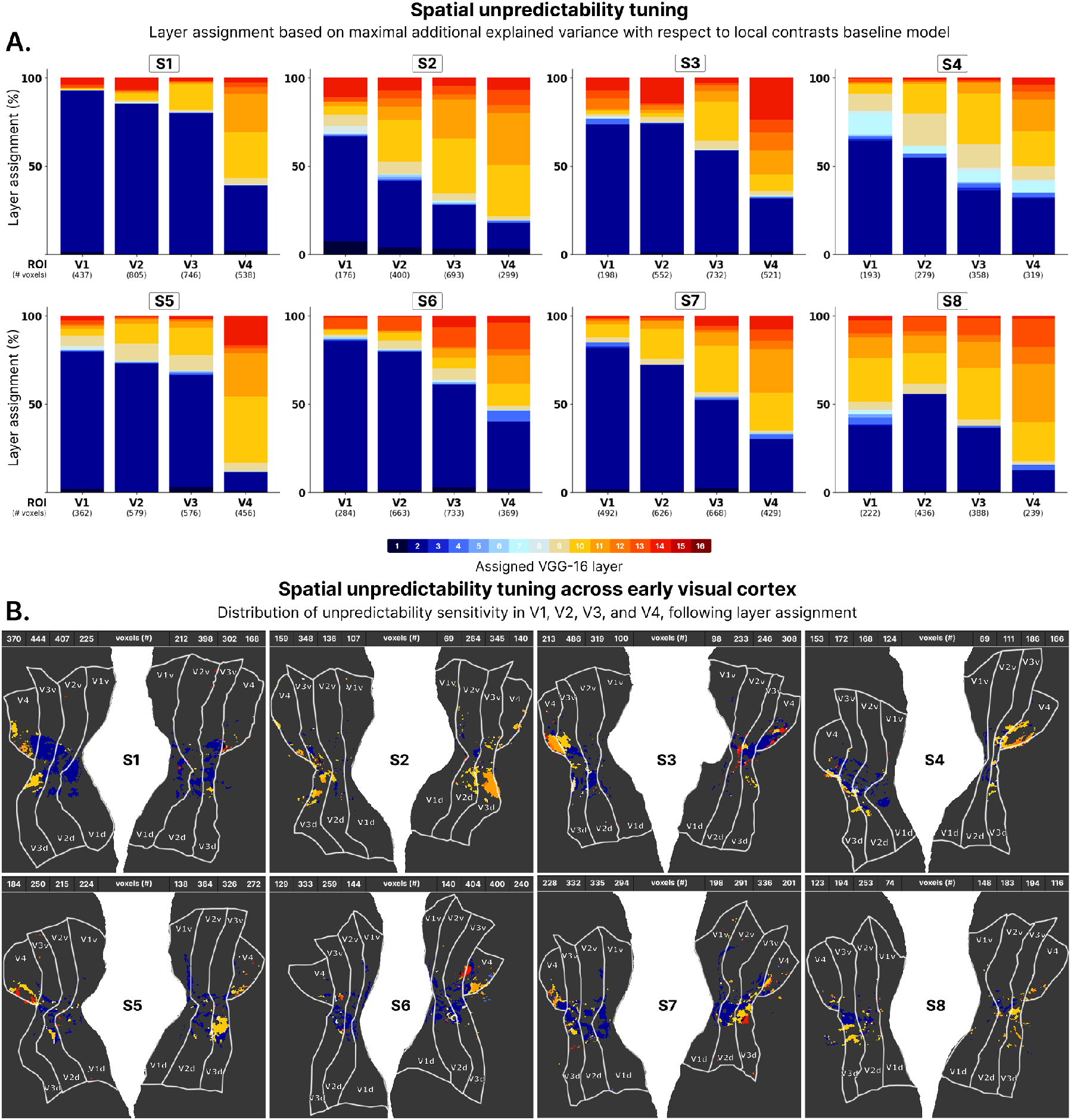
Consistent hierarchical tuning for spatial unpredictability across visual cortex. A) Voxel assignment based on the abstract level at which spatial unpredictability best explains neural variance, over and above the local contrast baseline model. Different levels of abstraction are represented by VGG-16 layers from which features were derived. B) Projection of spatial unpredictability effects on the flattened visual cortical surface of each subject, highlighting the gradual progression of higher-level sensitivity across the cortex.

**Figure S5:**
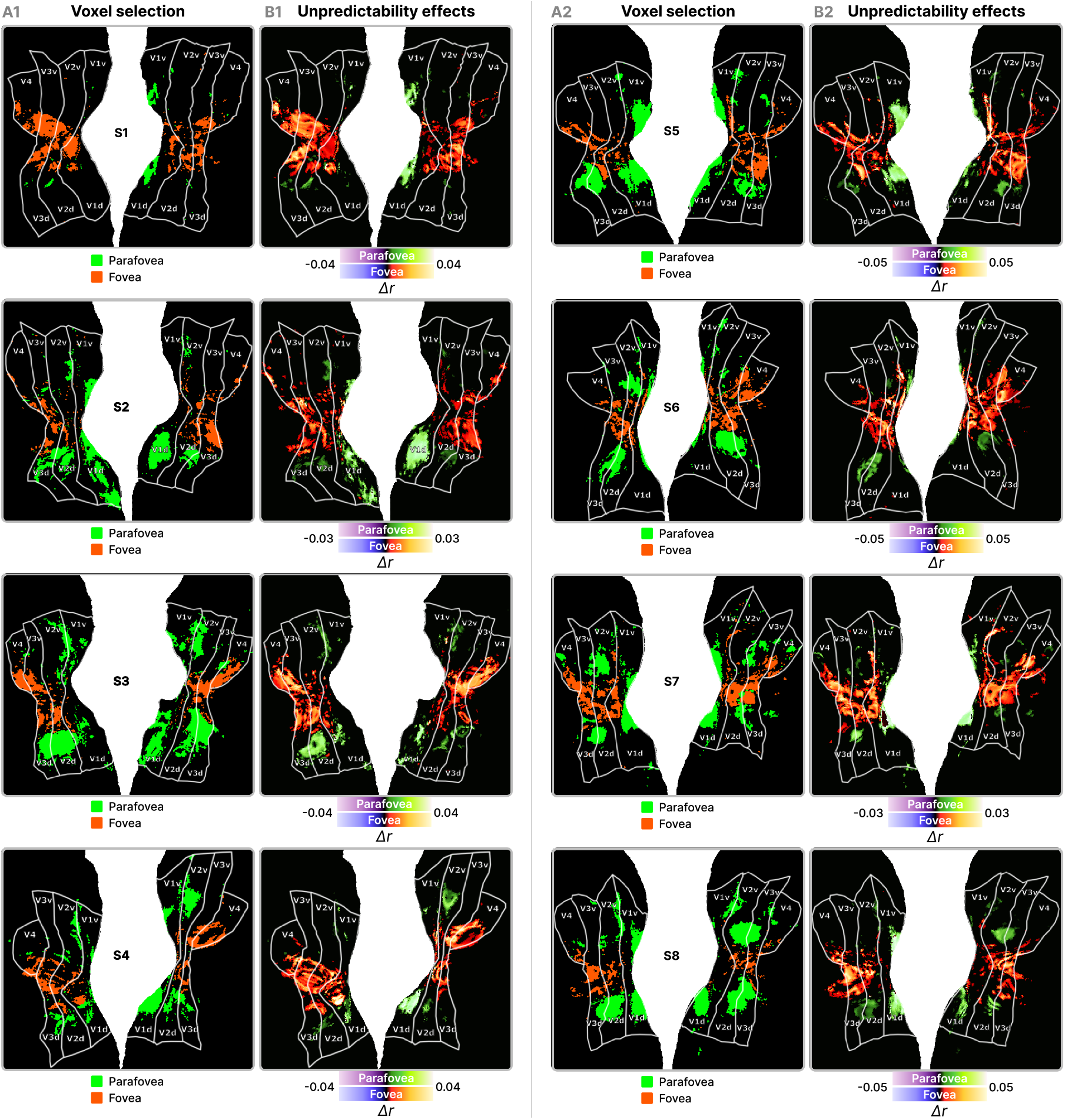
Stronger sensitivity to spatial unpredictability in more peripheral visual cortex than in central parts. A) Selection of voxels with either a pRF inside the central or one of the parafoveal image patches, subject-specific. B) Spatial unpredictability effects in the fovea and parafovea, as indicated by the additional explained neural variance (Δ*r*) over and above baseline model predictivity of patch-specific evoked responses.

**Figure S6:**
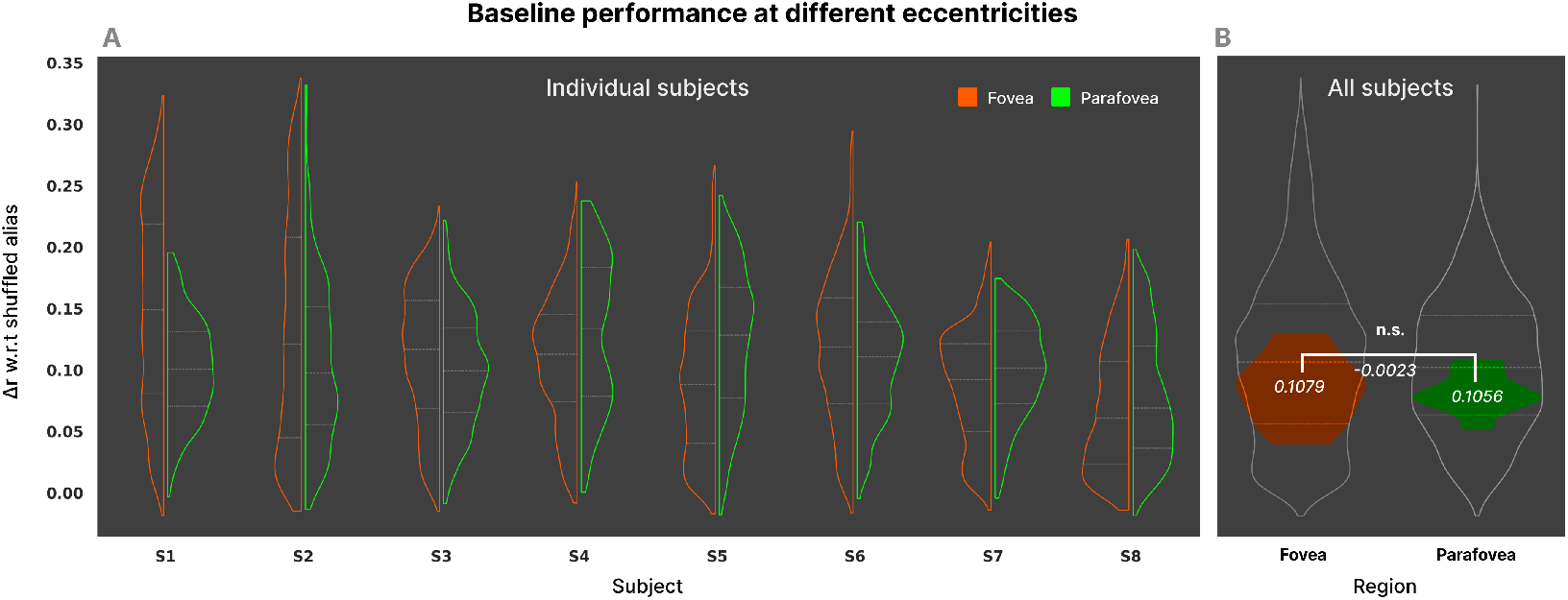
No consistent difference in neural predictivity of local-contrast baseline at different eccentricities. A) Neural predictivity of the local-contrast baseline model for both the foveal and parafoveal voxel selections in individual subjects, expressed as the difference in the Pearson’s *r* correlation of the model-predicted and actual neural response between the true baseline model and a shuffled version of it. B) Subject-aggregated baseline performance at different eccentricities. The gray outline shows the full voxel-specific data distribution across all subjects; filled colored distributions show subject-level means (fovea: mean Δ*r* versus shuffled control = 0.1079 *±* 0.0244; parafovea: mean Δ*r* versus shuffled control = 0.1056 ± 0.0148). A two-tailed bootstrapped t-test (10,000 iterations) on these 8 subject means per region revealed no significant difference (effect size = -0.0023, 95% CI [-0.0022, -0.0018], *p* = 0.8306).

**Figure S7:**
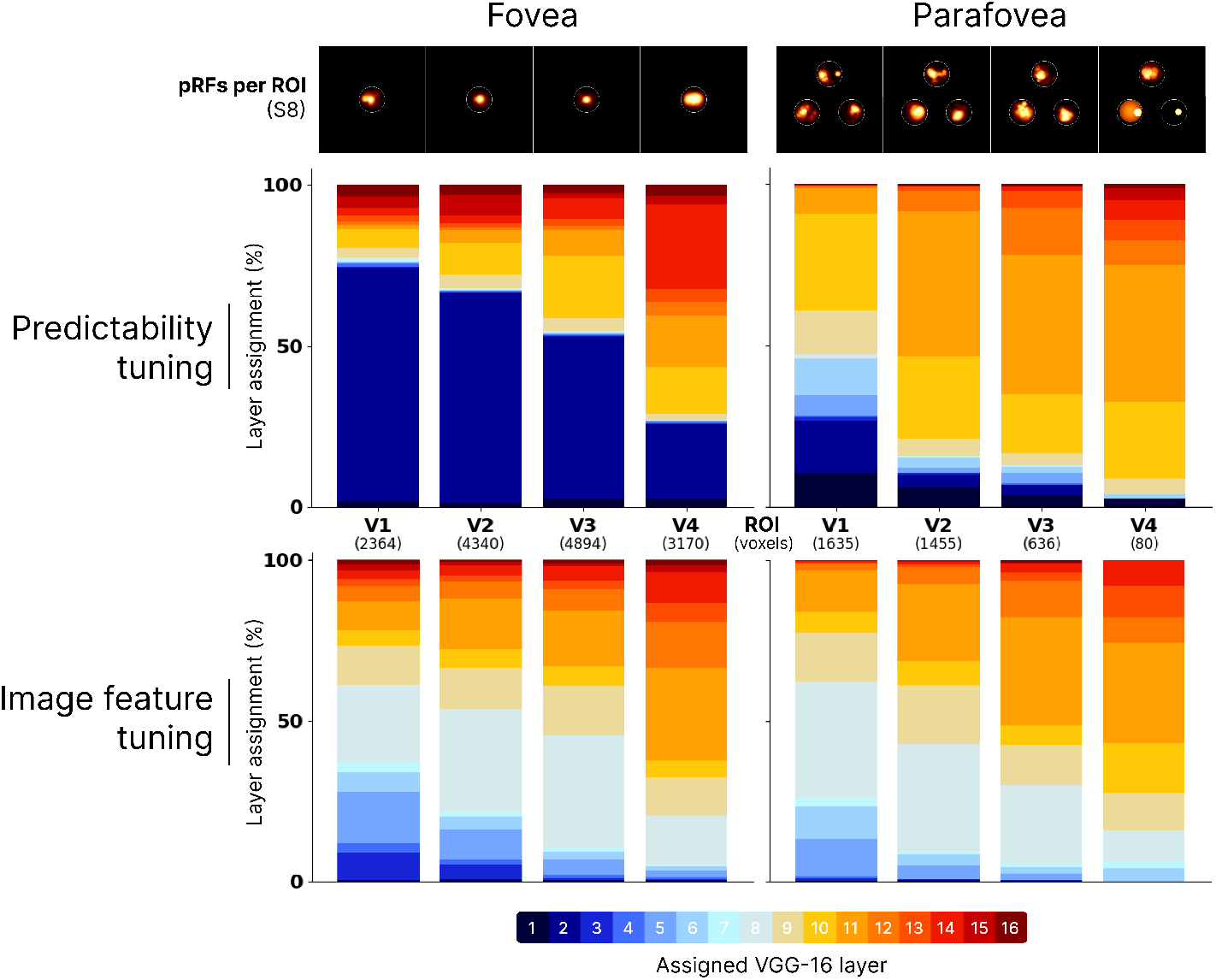
The parafovea is more tuned for high-level spatial unpredictability than the fovea, their bottom-up feature tuning is similar. Voxels with pRFs in parafoveal patch regions are more sensitive to spatial unpredictability at higher levels of abstraction than foveal voxels, indicated by the different voxel-wise layer assignments of the VGG-16 layer at which unpredictability features explain neural variance best (top row). Sensitivity to features encoded in the feedforward sweep of visual processing is more similar across the fovea and parafovea, as shown by the level and gradient of bottom-up encoded visual feature tuning (bottom row).

